# Assessing the impact of post-mortem damage and contamination on imputation performance in ancient DNA

**DOI:** 10.1101/2023.12.17.572049

**Authors:** Antonio Garrido Marques, Simone Rubinacci, Anna-Sapfo Malaspinas, Olivier Delaneau, Bárbara Sousa da Mota

**Affiliations:** Department of Computational Biology, University of Lausanne, Switzerland; Division of Genetics, Department of Medicine, Brigham and Women’s Hospital and Harvard Medical School, Boston, MA, USA; Program in Medical and Population Genetics, Broad Institute of MIT and Harvard, Cambridge, MA, USA; Swiss Institute of Bioinformatics, University of Lausanne, Switzerland; Regeneron Genetics Center, Tarrytown, New York, USA

## Abstract

Low-coverage imputation is becoming ever more present in ancient DNA (aDNA) studies. Imputation pipelines commonly used for present-day genomes have been shown to yield accurate results when applied to ancient genomes. However, *post-mortem* damage (PMD), in the form of C-to-T substitutions at the reads termini, and contamination with DNA from closely related species can potentially affect imputation performance in aDNA. In this study, we evaluated imputation performance i) when using a genotype caller designed for aDNA, ATLAS, compared to bcftools, and ii) when contamination is present. We evaluated imputation performance with principal component analyses (PCA) and by calculating imputation error rates. With a particular focus on differently imputed sites, we found that using ATLAS prior to imputation substantially improved imputed genotypes for a very damaged ancient genome (42% PMD). For the remaining genomes, ATLAS brought limited gains. Finally, to examine the effect of contamination on imputation, we added various amounts of reads from two present-day genomes to a previously downsampled high-coverage ancient genome. We observed that imputation accuracy drastically decreased for contamination rates above 5%. In conclusion, we recommend i) accounting for PMD by using a genotype caller such as ATLAS before imputing highly damaged genomes and ii) only imputing genomes containing up to 5% of contamination.

## Introduction

Over the past decade, there has been a fast increase of sequenced ancient genomes^1^. The growing amount of ancient genetic data allows for a better understanding of the evolutionary history of humans and other species, as shown by a multitude of discoveries in recent years^2–5^.

The field of ancient DNA (aDNA) research is accompanied by a unique set of challenges, namely, *post-mortem* damage (PMD) and contamination. PMD manifests as DNA fragmentation, and pronounced C-to-T deamination at the ends of DNA fragments^6^, that accumulate over time due to the absence of DNA repair mechanisms following an organism’s death. PMD can complicate sequencing and lead to erroneous genotype calls^7^. A strategy to mitigate C-to-T deamination is the application of a uracil-DNAglycosylase (UDG) and endonuclease VIII treatment during laboratory preparation^8,9^. UDG is an enzyme that recognizes and removes uracil residues arising from the deamination of cytosine^10^. The removal of uracil by UDG creates abasic sites, which are then recognized and cleaved by an endonuclease VIII. By actively targeting and removing uracil from aDNA, this treatment minimizes the incidence of post-mortem C-to-T deamination, thereby enhancing the accuracy of subsequent analyses.

Despite meticulous procedures intended to prevent contamination^11–13^, it is hard to avoid and sources of contamination can range from unintentionally introduced human DNA during sample extraction or laboratory processing^14–16^, to the persistent presence of DNA from microbial and environmental sources^17^. Microbial and environmental contamination are responsible for the underrepresentation of endogenous DNA. The competitive amplification of this non-target DNA results in the ancient DNA of interest to become overshadowed and to numerous genomic positions being either weakly covered or entirely devoid of sequencing reads. While we can practically remove all microbial and environmental contamination by mapping the sequenced reads, contamination with DNA from the same or closely-related species cannot be fully eliminated. Even at minimal levels, this type of contamination can skew downstream analyses, leading to potentially erroneous conclusions. As such, contamination estimation is a standard quality control step in aDNA studies. Genomes with a contamination level exceeding 5% (an adhoc threshold used in several studies) are often discarded, resulting in the loss of potentially informative data.

Both PMD and contamination contribute to reduced genome coverage. On the one hand, PMD produces shorter DNA fragments with reduced mappability, and the deaminations introduced by PMD can increase discrepancies relative to the reference genome, making the fragments harder to map^18,19^. On the other hand, microbial contaminant DNA tends to be several times more abundant than the endogenous DNA that is under study, resulting in low amounts of the latter and, consequently, low depth of coverage.

A standard approach to address the challenge of low coverage in ancient genomes is pseudo haploidization. This method involves selecting one single read to represent a haploid genotype at each genomic position^13^. However, this method carries inherent risks such as under-estimation of allele frequencies and may inadvertently introduce a bias towards the reference genome^20–22^.

A different approach is genotype imputation. This is a method that makes use of statistical models to infer missing genotypes in a target sample, guided by a reference panel of haplotypes^23^. In the case of SNP-array data, the process begins with the phasing of the target sample to determine the underlying haplotype structure. Then, the method infers missing genotypes by identifying stretches of shared haplotypes between the target sample and the reference panel^24^. Given the nature of ancient genomes, one should apply low-coverage WGS (whole-genome sequencing) imputation methods. In this case, genotype likelihoods (GLs) are the input data for imputation. These GLs are the probabilities of finding certain genotypes based on the available sequencing data, and can thus encode the uncertainty inherent to low-coverage ancient genomes. Unlike SNP-array imputation, in low-coverage WGS imputation the phasing and imputation steps happen simultaneously^25–27^. Moreover, it has been shown that low-coverage WGS imputation can be more informative than SNP-array imputation^28^.

While primarily used for present-day low-coverage genomes, recent studies have demonstrated the effectiveness of genotype imputation of ancient genomes^29–31^. Nevertheless, while this method addresses the low-coverage problem in aDNA, it is unclear how contamination and PMD affect imputation and whether we could use imputation to tackle these challenges as well. Although it was shown that imputation can have a corrective effect on deaminated sites^31^, it is not clear to what extent PMD affects imputation performance, and hence what the best practices are regarding pre-imputation PMD filtering. Moreover, to our knowledge, there has been no research on the impact of contamination on imputation. Thus, a description of imputation impact on contaminated sites and vice-versa is lacking.

In this study, we aim to i) assess whether imputation performance can be improved by using a genotype caller specifically designed to account for PMD and ii) study the impact of imputation on contamination levels (i.e., can genotype imputation “correct” contaminated positions so as to retrieve the original genotypes?). To address these aims, we performed simulations to mimic the properties of ancient DNA. We achieved this by downsampling high-coverage ancient genomes to low coverage, and then performing imputation experiments. When investigating the effect of PMD, we compared two imputation pipelines: a combination of GLIMPSE with GLs generated with either i) bcftools^32^, a widely used genotype caller, or ii) ATLAS^33^, a genotype caller designed to explicitly take into account PMD when generating the calls. To determine how contamination affects imputation, we artificially contaminated an ancient genome with varying amounts of sequencing reads coming from present-day genomes. In the following sections, we show that, while PMD filtering has a noticeable impact in the presence of high deamination rates, it does not substantially improve imputation performance in other cases. Furthermore, we demonstrate that contamination can drastically reduce imputation accuracy, rendering imputed genotypes unreliable.

## Results

### Methodology for comparing imputation accuracy of ancient genomes using distinct genotype callers

To assess the impact of PMD on imputation accuracy, we imputed two sets of GLs that we called using either i) ATLAS or ii) bcftools, and compared these to validation datasets through non-reference discordance (NRD) estimates and principal component analysis (PCA), focusing on SNPs differently imputed across the two datasets. ATLAS, specifically designed for aDNA, infers PMD patterns and recalibrates base quality scores. In contrast, bcftools serves as a conventional genotype caller providing a comparative dataset. We performed these imputation experiments using seven high-coverage ancient genomes of diverse origins and varying PMD rates (**Table S1, Supplementary Figure 1** and **Supplementary Figure 2**). We downsampled these genomes to emulate low-coverage (1x) conditions, and used the high-coverage genomes as validation (**Figure 1A**). We generated three validation datasets: i) by calling genotypes with ATLAS (“ATLAS validation”), ii) by calling genotypes with bcftools (“bcftools validation”), and iii) by intersecting the two previous genotype sets so as to only keep concordant sites between the two (“validation concordant”). However, we mostly relied on the latter due to inherent difficulties in establishing a ground truth for ancient genomes (**Supplementary Figure 3**). Afterwards, we performed genotype imputation on the two computed sets of genotypes likelihoods using GLIMPSE v1.1.1^27^, a genotype imputation tool for low-coverage whole-genome sequencing data. To evaluate imputation performance, we compared our results with the three different validation datasets using the NRD metric, defined as NRD = (eRR + eRA + eAA) / (eRR + eRA + eAA + mRA + mAA), where ‘e’ and ‘m’ represent the number of errors and matches, respectively, while ‘R’ and ‘A’ stand for reference and alternative alleles, respectively. We calculated NRD on all the 1000 Genomes bi-allelic SNPs^34^, on the intersection of the 1000 Genomes and 1240K bi-allelic SNPs^35–37^, and on a set of 2.8M SNPs (see Methods section). Restricting to the 2.8M SNPs, we also performed PCA using smartPCA^38,39^ to visualize and measure where imputed samples are positioned relative to their validation concordant set across the first 10 principal components (PCs). The PCA components were calculated using worldwide present-day genomes from SGDP^40^ and we projected the ancient data onto these. For each ancient individual, we projected four datasets: i) validation concordant, ii) identically imputed genotypes when using the two GLs sets (“imputed concordant”), and the differently imputed genotypes across the two GLs datasets computed with iii) ATLAS (“ATLAS imputed discordant”) and iv) bcftools (“bcftools imputed discordant”). Finally, for each individual sample, we calculated the Euclidean distance to their validation and adjusted by the corresponding eigenvalue of each PC with a focus on the subset of SNPs that are discordant.

**Figure 1:**
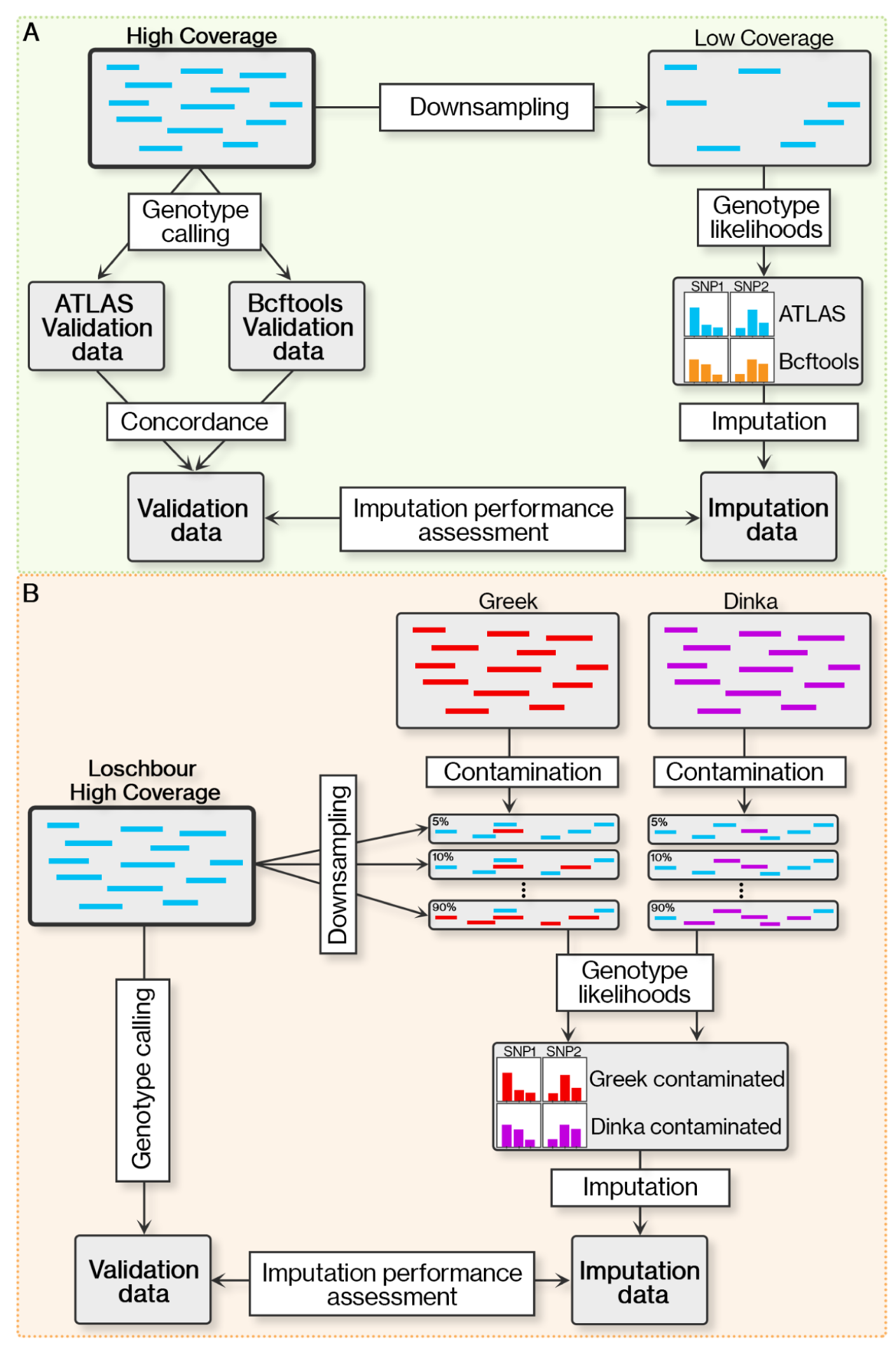
Schematic representation of the methodology. **A)** Outline of the approach used to evaluate the effect of two different genotype callers on the imputation accuracy of ancient genomes. **B)** Methodology followed to gauge the impact of contamination on genotype imputation of ancient genomes.

### Memory and time requirements of ATLAS and bcftools to generate genotype likelihoods

To assess the computational efficiency of genotype calling with either ATLAS or bcftools, we measured running times and memory usage when computing GLs for the 1x downsampled and the high-coverage genomes (**Supplementary Figure 4**). We observed that ATLAS required considerably longer running times. For chromosome 1 and the downsampled genomes, ATLAS took approximately 48 times more time than bcftools, averaging ∼1.6h per sample, while for the high-coverage genomes, ATLAS took ∼5.9h, that is, 17 times more time than bcftools. Genome-wide, ATLAS also required substantially more memory resources, using around 45 times the memory allocation of bcftools with an average of ∼182 Gb per sample for the downsampled genomes, and ∼3,200 Gb for the high-coverage genomes (768 times more memory than bcftools for the same genomes). It should be noted, however, that generating GLs with ATLAS entails three steps instead of one, as is the case of bcftools: i) splitting of single-end read groups according to read length, and merging of paired-end reads if present, ii) generating empirical estimates for PMD, which are then used to refine the calling step, and iii) genotype calling.

### Impact of taking PMD into account on imputation accuracy

#### Accounting for PMD improves imputation performance in highly damaged individual samples

When restricting to identically imputed genotypes using ATLAS and bcftools GLs, that is, the imputed concordant set, we observed that these data were placed on the PCA space close to present-day individuals from the same continents, as expected. Furthermore, these datasets co-localised with their respective validation (validation concordant) (**Figure 2A**). We found that the imputed ATLAS discordant data was closer to the validation for all individual samples except NE1^41^ (9.6% PMD), as depicted in **Figure 2A**,**B**. Different trends were observed when using the 1240K SNPs (**Supplementary Figure 5**), though. The biggest differences between imputed ATLAS and bcftools data were found for BOT2016^42^ (20.8% PMD, 0.51% discordant sites), Sumidouro5^43^ (42.0% PMD, 0.86% discordant sites), Yana^44^ (13.9% PMD, 0.92% discordant sites), and ela01^45^ (3.0% PMD, 1.67% discordant sites). This disparity is particularly noticeable for Sumidouro5, that exhibits the highest PMD rate (42.0%), and whose discordant imputed ATLAS genotypes are considerably closer to the validation. In the case of Yamnaya^42^ (18.8% PMD, 0.26% discordance sites), the two discordant datasets were similarly close to their validation (weighted Euclidean distances of 0.40 vs. 0.42 for ATLAS and bcftools discordant data, respectively). In conclusion, the discordantly imputed bcftools GLs tend to be more different from the validation, and thus less accurate than the imputed ATLAS GLs. However, the proportions of the discordantly imputed SNPs were relatively low, ranging between 0.26% and 1.67%, indicating a high level of similarity between the two imputed datasets.

**Figure 2:**
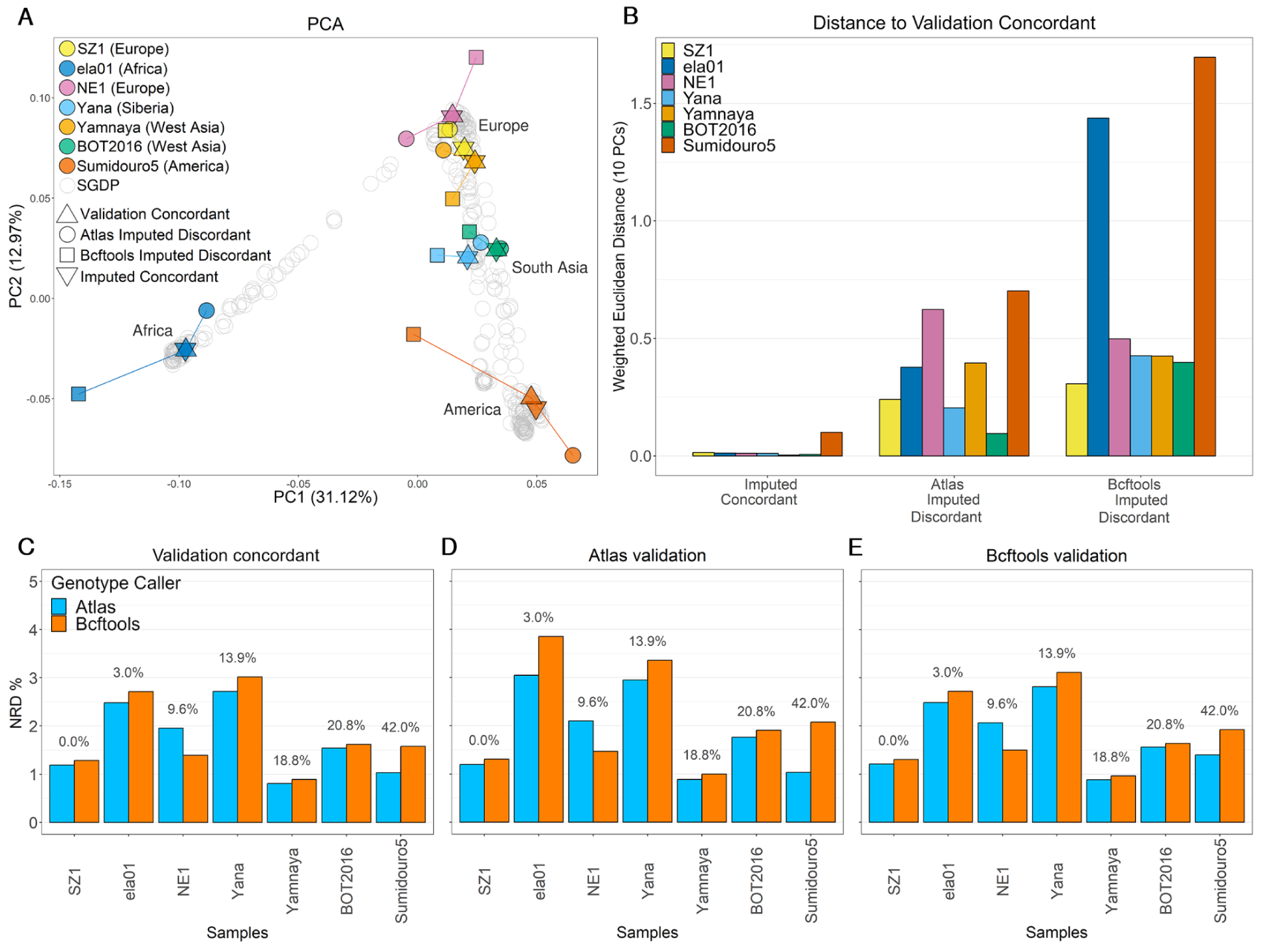
Effect of two different genotype callers on imputation accuracy. **A)** Two first principal components of principal component analysis (PCA) of present-day genomes (SGDP). The genetic data of seven ancient individuals were projected onto it. For each individual, the triangle shape indicates the validation concordant dataset, the inverse triangle the set of imputed positions in agreement with both tools, and the circles and square shapes the positions differently identified by ATLAS (“ATLAS Imputed Discordant”) and bcftools (“Bcftools Imputed Discordant”), respectively. The continental origins of the individual samples are indicated on the legend. **B)** Weighted Euclidean distances for the seven ancient individuals. We calculated the Euclidean distances between the validation concordant set of each sample, and their concordant and discordant imputed genotypes. We measured the distances across the first 10 PCs, and subsequently weighted the distances by the eigenvalue of each PC. **C)** Non-reference discordance (NRD) for the seven ancient individual samples when called with either ATLAS (blue) or bcftools (orange) prior to imputation. The values on top of each sample indicate the PMD rate, and samples were ordered by increasing PMD rates. NRD was assessed using three validation datasets: (**C**) validation concordant, (**D**) ATLAS validation and (**E**) bcftools. For NRD assessment on all variants, see **Supplementary Figure 5F-H**.

An examination of the imputation error rates allows for a deeper understanding of how imputation is affected by distinct genotype callers. We computed NRD and other error rates using three validation sets obtained from the high-coverage genotypes: i) bcftools calls, ii) ATLAS calls and iii) validation concordant (intersection of i) and ii)). We computed NRD for three different sets of SNPs: i) the same SNPs as in the abovementioned PCA analysis (2.8M SNPs) (**Figure 2C-E**, for more detailed error rates see **Table S2**), ii) the intersection of the 1000 Genomes and 1240K bi-allelic SNPs all sites (**Supplementary Figure 5C-E**), and iii) all sites in the 1000 Genomes panel (**Supplementary Figure 5F-H**). At first sight, the (2.8M SNPs) NRD results corroborate the previous observations regardless of the validation dataset. Imputation using ATLAS GLs resulted in lower NRD for ela01, Yana and Sumidouro5, while the opposite was seen for NE1. We caution that the NRD values calculated at all 1000 Genomes SNPs yielded the opposite trend for ela01. For SZ1^46^, Yamnaya and BOT2016, the two imputed datasets had approximately the same NRD values, ∼1.23%, ∼0.85% and ∼1.58%, respectively. While such a result was expected for SZ1, given that its libraries were UDG-treated and hence no gain in accuracy was expected with ATLAS, PMD is prevalent in the Yamnaya and the BOT2016 genomes (18.8% and 20.8% PMD, respectively) and we would expect ATLAS to have a considerable impact. Therefore, using one or the other genotype caller prior to imputation did not systematically affect imputation performance given PMD rates as expected. This suggests that there are other factors affecting GLs computation and/or imputation that ultimately impact imputation performance in a non-trivial way. Nonetheless, Sumidouro5 represents the clearest example of how using ATLAS to generate GLs prior to imputation can result in higher imputation accuracy, which is particularly visible when using the ATLAS validation dataset as the ground truth. In this case, imputed ATLAS GLs yielded 1.03% NRD, two times smaller than the NRD for imputed bcftools GLs (2.08%).

#### Transition substitutions proportions tend to increase in discordantly imputed variants

To determine whether the differences between the two imputed datasets arose from the fact that ATLAS takes into account PMD upon calculating GLs, we examined whether there was an overrepresentation of transitions, i.e., affected by PMD, among the discordantly imputed sites. The transition base change proportion in the imputation reference panel was 68.7% (**Table S3**). In comparison, the average transition proportion for concordantly imputed positions across all samples aligned with this expectation. At discordantly imputed positions, we found the proportion of transition sites to be similar to the overall transition rate for Yamnaya, NE1 and BOT2016. There was a clear increase in transition representation among discordantly imputed SNPs for Yana (71.1%) and Sumidouro5 (73.0%). Even though there was no clear correlation between PMD and transition numbers increase, the largest difference was again found for Sumidouro5. For this genome, there were 2.7 times more transitions than transversions among the discordantly imputed sites, compared to 2.2 times in all considered sites, further suggesting that ATLAS aids imputation in the presence of very high PMD rates. We observed similar trends of transition base change proportions when considering the intersection of the 1000 Genomes and 1240K SNPs, and the 2.8M SNPs used in PCA analyses (**Table S4** and **Table S5**).

#### Inherent difficulties in assessing imputation performance of African individual samples

While Sumidouro5 and ela01 have the largest discrepancies between their imputed datasets, the reasons underlying those differences are clearly distinct, given that ela01 has a much lower PMD rate (3%) and thus we would not expect imputation to considerably benefit from ATLAS GLs. Specifically, ela01’s performance with ATLAS also stands out as shown by its Euclidean distance to the ground truth of 0.38, in contrast to 1.44 with bcftools (**Figure 2B**), and reduced NRD values for ATLAS (2.48%) compared to bcftools (2.71%) with the validation concordant (**Figure 2C**). However, the NRD calculated on all 1000 Genomes positions using the validation concordant showed a different trend. Here, bcftools’s imputed GLs yielded a lower NRD of 4.06%, compared to 4.61% with ATLAS (**Supplementary Figure 5F-H**). The distinct characteristics of ela01 are further highlighted by its elevated discordance rate of 1.67% at the 2.8 million SNPs, nearly double that of most other samples (**Table S6**). Additionally, ela01 exhibits among the highest imputation error rates overall, irrespective of the genotype caller used (**Table S2**). Finally, ela01 stands out as having the lowest proportion of transition base changes in the discordant positions when considering all imputed SNPs (66.7% compared to 68.7% for all sites, **Table S3**). These findings suggest that evaluating imputation performance in African samples may be challenging due to the unique features and complexities of their genomic profiles and their ancestries being underrepresented in most reference panels.

### Assessing how contamination affects imputation performance

#### Methodology for assessing imputation’s ability to rectify contamination in ancient genomes

To assess the potential of genotype imputation in mitigating contamination, we virtually contaminated the Loschbour genome, an 8000-year-old Western Hunter-Gatherer^47^, with present-day human DNA from a European Greek and an African Dinka genomes from SGDP^40^, as depicted in **Figure 1B**. To ensure an accurate representation of the impact of contamination on a typical ancient sample, the Loschbour genome was initially downsampled to low coverage (1x). Then, based on the desired contamination levels ranging from 5% to 90%, we substituted a proportionate number of reads - drawn at random - from the ancient genome with reads from the modern genomes. We verified that the genome was contaminated as expected by estimating contamination on the X chromosome using contaminationX^48^ (**Supplementary Figure 6**). Afterwards, we computed the genotype likelihoods on the two artificially-contaminated samples using bcftools, and imputed them with GLIMPSE. In order to assess to which degree imputation rectified contamination and allowed the retrieval of the high-coverage genotypes, we compared the contaminated samples with the original non-contaminated high-coverage Loschbour by calculating NRD and evaluating their relative positions on PCA whose PCs were calculated with present-day genomes. Finally, we verified whether contamination has a different signature at the haplotype level when compared to admixture. For that, we inferred local ancestry for imputed genomes that had been previously contaminated with Dinka DNA using three reference populations from the 1000 Genomes panel: CEU (Utah residents with Northern and Western European ancestry), LWK (Luhya in Webuye, Kenya) and YRI (Yoruba in Ibadan, Nigeria).

#### Imputation severely impacted by contamination

We projected the non-contaminated high-coverage Loschbour genome, the two present-day Greek and Dinka genomes, and the imputed 1x contaminated genomes onto the PCA built with the SGDP dataset to determine how far the latter were relative to the non-contaminated Lochbour genome. We observed that, as contamination increased, the imputed sample moved further away from the non-contaminated Loschbour (**Figure 3A**). Particularly, in the case of Dinka-contaminated samples, it appears that, with an increasing contamination rate, the samples moved progressively closer to the genetic signature of the original Dinka individual. With increasing contamination by present-day Greek DNA, the samples similarly deviated further from the Loschbour original sample, although in a less linear trajectory. However, this trend became linear when restricting the analysis to only the Western Eurasian populations in SGDP (**Figure 3B**), as well as at higher PCs (PC7 vs. PC8 in **Supplementary Figure 7**). The observed nonlinearity may be due to these dimensions not distinctly separating the Loschbour and Greek populations. In contrast, the Dinka contamination displays a linear trend as PC1 primarily distinguishes African versus non-African populations^49^. In addition, we verified the effect of contamination rate on the number of heterozygous sites. As expected, a maximum number of heterozygous sites was reached at around 50%-contamination rate, that is, 7.5% and 6.4% for the Dinka and the Greek contaminants, respectively, in comparison with 3.4% at 0% (imputed uncontaminated 1x Loschbour genome) and 5.7% (Dinka) and 4.5% (Greek) at 100% contamination rates, respectively (**Supplementary Figure 8**).

**Figure 3:**
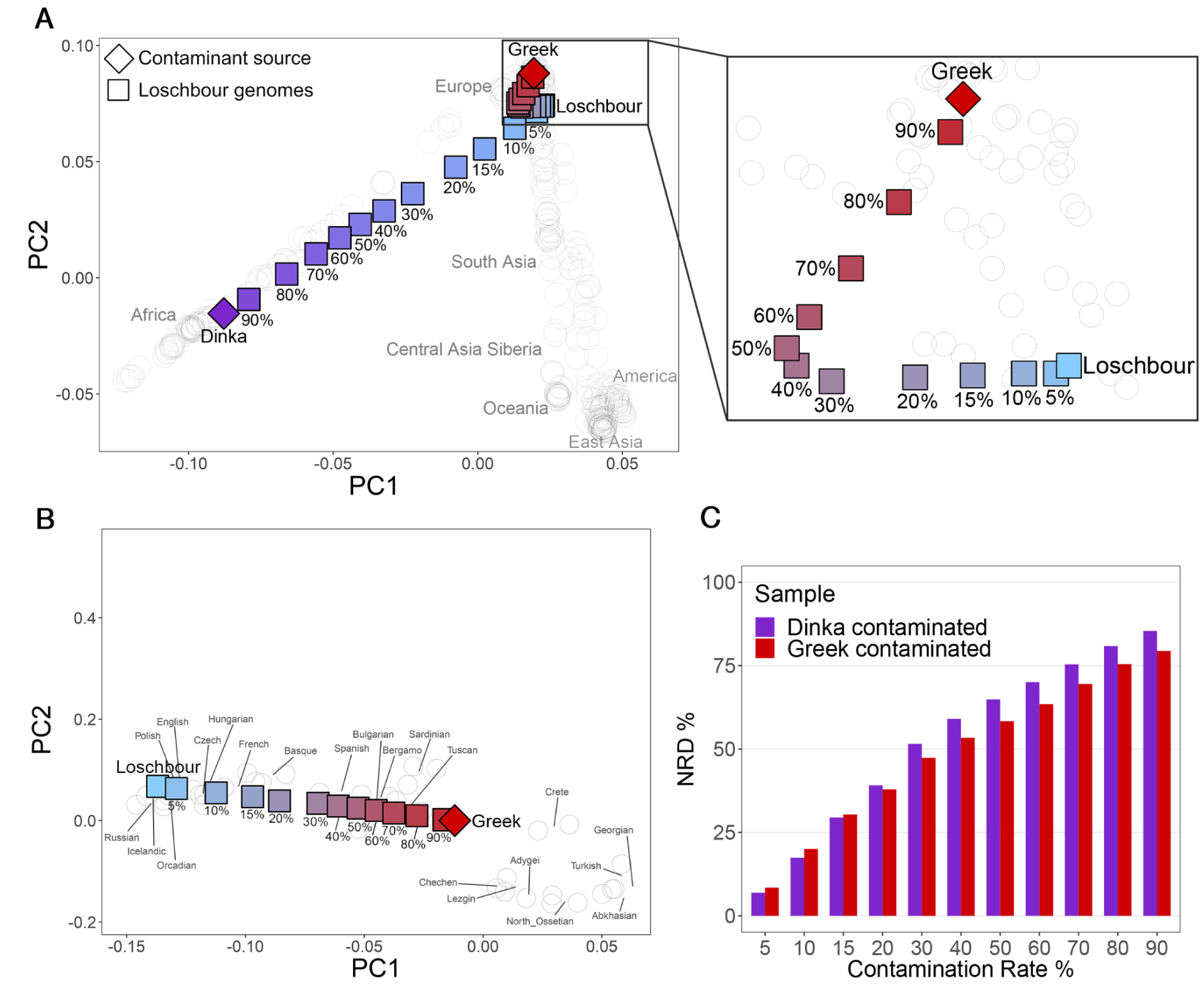
Effect of contamination on genotype imputation. Principal component analysis (PCA) of present-day genomes and projection of uncontaminated and imputed contaminated ancient genomes. The ancient downsampled (1x) Loschbour genome (light blue) was subjected to varying degrees of contamination, ranging from 5% to 90%, with DNA from a present-day Greek individual (red) and a present-day Dinka individual (purple). **A)** PCA of worldwide present-day populations (SGDP), represented by empty gray circles, and projection of all the imputed contaminated downsampled Loschbour genomes. **B)** PCA of present-day Western Eurasians, with a focus on the Greek-DNA imputed contaminated Loschbour genomes. **C)** Non-reference discordance (NRD) as a function of contamination levels, having the high-coverage Loschbour genome as ground truth. Red and purple bars indicate Greek-contaminated and Dinka-contaminated Loschbour samples, respectively.

Imputation of the uncontaminated Loschbour genome downsampled to 1x resulted in an NRD of 2.72% (**Figure 3C**). In the presence of 5% present-day Greek DNA contamination, NRD increased to 8.4%, which became 79.4% at 90% contamination. Similarly, samples contaminated with present-day Dinka DNA had an NRD of 6.9% at 5% and 85.4% at 90% contamination. The implications are that, as contamination levels rise, the reliability of imputed genotypes for downstream analyses deteriorates. Moreover, contamination from present-day Dinka DNA seemed to have a more important detrimental impact on imputation accuracy relative to contamination from present-day Greek DNA. Specifically, beyond a contamination level of 20%, the NRD associated with Dinka contamination exceeded that of its Greek counterpart. Furthermore, squared Pearson correlation values (**Supplementary Figure 9**) highlighted that Greek-contaminated samples outperformed Dinka-contaminated ones in imputation accuracy. This observation suggests that imputation performance was more negatively affected by higher levels of contamination coming from a more distantly related source. These results indicate that contamination substantially compromises genotype imputation accuracy, making it ineffective to rectify contamination in ancient genomes. We hypothesize that contaminant reads prevent the imputation model from copying from the most informative reference haplotypes, as contaminated genomes contain information from more than two haplotypes (four, in the case of a single contamination source).

#### Contamination leaves a distinct signal at the haplotype level

Admixture and contamination could leave similar genetic hallmarks, as implied by the findings that i) similarity between imputed contaminated genomes and their contamination sources increased with contamination rate and ii) heterozygosity peaked at 50% contamination. When inferring local ancestry for imputed genomes contaminated with Dinka DNA, we found that haplotype tracts with an African origin were only inferred for contamination rates above 15%, while true admixed tracts can be found at much lower admixture levels (**Supplementary Figure 10A**,**B**). Furthermore, at 50% contamination rate, for instance, the tract length distribution was different from the expected distribution for 50% admixture levels, as exemplified with local ancestry inferred in an African American individual (**Supplementary Figure 10C**). In contrast to true admixture, we found an enrichment for short tracts across the genome (**Supplementary Figure 10D**,**E**), reflecting that contamination affects all sites in an uniform fashion and not according to a recombination map as in the case of admixture.

## Discussion

Genotype imputation has been increasingly employed in aDNA studies, unlocking downstream applications that were typically limited to high-coverage genomes, thus allowing to tackle evermore questions. It is therefore essential to thoroughly understand its limitations and potential. Our study provides insights into how PMD, a common feature of ancient genomes, and contamination affect genotype imputation of ancient genomes. Firstly, we found that taking PMD into account prior to imputation yielded unequivocally more accurate genotypes in the presence of very high PMD rates. Although imputation of ATLAS GLs led to overall lower imputation error rates, these constituted modest gains for most of the remaining genomes regardless of their PMD rates. This result suggests that imputation is robust to some level of damage. As such, ATLAS’s high computational needs and extended run times may not justify its benefits when ancient genomes are not highly damaged. Secondly, we showed that genomes with more than 5% contamination are inaccurately imputed. This is also the current threshold used to discard contaminated samples in aDNA studies. Moreover, imputation accuracy decreased with the increase of contamination levels in such a way that the imputed contaminated genomes were increasingly closer to contamination sources. Nonetheless, the contamination signature was clearly distinct from admixture at the haplotype level. This observation suggests that it is possible to detect and estimate nuclear DNA contamination using a framework built upon imputation and by exploiting disruption of linkage disequilibrium by contamination as in Nakatsuka et al^50^. Another potential future methodological development could be aimed at correcting PMD in ancient genome imputation, since we have shown before that imputation can correct deaminated sites. A full description of how and to what extent this correction works is however needed to that end. It also remains to be assessed how imputation varies across the genome, which can have implications for studies focused on particular genome regions, such as selection studies. Finally, a limitation of imputation not restricted to ancient genomes is the fact that present-day imputation reference panels do not contain private variation from under/non-represented populations. Such limitations can be mitigated by using more diverse reference panels and/or by assembling reference panels that include ancient genomes so as to recover variation that has been lost over time. However, high-coverage ancient genomes are the exception and their sample size is probably too small to improve imputation performance at private variants.

## Methods

### a) Datasets

#### i) Ancient genomes

We collected eight high-coverage ancient genomes from publicly available studies and reported them in **Table S1**. This table provides additional details about the origin of each sample, their coverage, and the average rate of PMD at the 5’ read ends. These samples offer a global representation and display varying damage rates. Certain samples, as indicated in **Table S1**, underwent a UDG-treatment to reduce PMD. The PMD values for the remaining samples were either stated as reported in the original study, or assessed by us. We selected seven ancient genomes for the first part of our experiment focusing on assessing how accounting for PMD impacts imputation accuracy, while the remaining genome, Loschbour, was used in the second part investigating the impact of contamination on imputation.

#### ii) Simons Genome Diversity Project

We used the Simons Genome Diversity Project (SGDP)^40^ dataset as a reference panel to perform PCA. This dataset includes genomic information from 300 individuals that come from 142 populations worldwide. In addition, we sampled reads from two SGDP genomes (a Dinka (B_Dinka-3) and a Greek individual (S_Greek-1)) to contaminate the Loschbour ancient genome.

#### iii) 1000 Genomes UMICH (imputation reference panel)

We used a version of the 1000 Genomes v5 phase 3 panel^34^ containing 2504 genomes resequenced at 30x, that was phased using TOPMed^51^, excluding singletons and only keeping biallelic sites. Furthermore, we lifted over the reference panel from b38 to hg19 using Picard liftoverVCF (https://gatk.broadinstitute.org/hc/en-us/articles/360037060932-LiftoverVcf-Picard-), with hg38ToHg19 chain from the University of California Santa Cruz liftOver tool (http://hgdownload.cse.ucsc.edu/goldenpath/hg38/liftOver/).

### b) PMD Assessment

We used the bamdamage tool from bammds^52^ to estimate PMD as the rate of C-to-T misincorporations at the 5’ end first position. This tool enabled us to evaluate the PMD rate for 6 of the 7 samples. For the remaining sample (SZ1), we relied on the damage report from the published study (**Table S1**).

### c) Downsampling

We downsampled all individual samples used in the study to 1x coverage using samtools^32^ v1.10.

### d) Genotype calling

We used bcftools^32^ v1.10 and ATLAS^33^ v0.9.9 to call genotypes at the 1000 Genomes bi-allelic SNPs and generate subsequent genotype likelihoods from the downsampled and high-coverage genomes, using the human reference genome humanG1Kv37 (https://www.ncbi.nlm.nih.gov/assembly/GCF_000001405.13/).

#### i) Bcftools

To generate genotype likelihoods from the downsampled genomes via bcftools, we used the *bcftools mpileup* command with parameters *-I, -E*, and *-a ‘FORMAT/DP’* with the *--ignore-RG* flag enabled, followed by the *bcftools call* command with the *-Aim* and *-C* alleles parameters.

Before calling genotypes from the non-UDG treated high-coverage genomes using bcftools, we trimmed 10 base pairs from each end of the reads using BamUtil^53^ v1.0.14 to reduce the impact of PMD. Then, we performed five additional quality control parameters:

1. We excluded positions presenting a minimum mapping quality of 30 (*-q 30*) and a minimum base quality of 20 (*-Q 20*), as well as reduced mapping quality for reads featuring an abundance of mismatches with the parameter *-C 50*.
2. We removed sites not included in the 1000 Genomes accessible genome strict mask^54^.
3. We filtered out sites located in regions known to contain repeat elements^55^.
4. We eliminated positions exhibiting extreme values of coverage depth. For the lower threshold, positions with depth less than one-third of the mean or less than eight (if the one-third mean coverage fell below eight) were filtered out. For the upper threshold, positions exceeding twice the mean depth of coverage were also excluded.
5. Finally, positions with a QUAL value below 30 (*QUAL<30*) were also omitted.

#### ii) ATLAS

To call the genotypes with ATLAS, we followed the standard pipeline for the analysis of individual samples as described in https://bitbucket.org/wegmannlab/atlas/wiki/Home. For the *splitmerge* step necessary to split single-end reads groups according to length and merge paired-end reads if present, we needed to divide samples containing mixed libraries, meaning, libraries presenting both single- and paired-end reads, into single-end and paired-end only libraries. To do that, for all mixed libraries we extracted the paired-end reads with *samtools view -f 1 -F 0* and the single-end reads with *samtools view -f 0 -F 1* into two separate bam files, then, we recombined all the libraries into an reunified bam file. This step was necessary to remove any mixed library and circumvent a problem related to mixed libraries that we encountered with the version of ATLAS we were working with. Afterwards, we used BAMdiagnostics in order to determine the maximum read length to give as input to *splitmerge*, estimated the empirical post-mortem damage with *PMD*, and used the *MLE* caller to generate genotype likelihoods. We applied this process to evaluate genotype likelihoods for both downsampled and high-coverage genomes.

Due to ATLAS not listing positions that are not covered by any read, we manually added those missing positions. This was a necessary step, because GLIMPSE v1.1.1 does not impute unlisted positions even if they are present in the imputation reference panel.

### e) Genotype Imputation

To impute the downsampled genomes, we used GLIMPSE v1.1.1^27^ as follows:

1. We used *GLIMPSE_chunk* to segment chromosomes into chunks ranging between 1-2 Mb in size, with an additional buffer region of 200kb flanking each chunk.
2. We performed imputation on the chunks using *GLIMPSE_phase* with the default parameters and using the 1000 Genomes panel as a reference panel.
3. Finally, we used *GLIMPSE_ligate* to ligate the imputed chunks into complete chromosomes.

### f) Imputation performance evaluation

In order to evaluate the imputation accuracy in both experiments, we used *GLIMPSE_concordance* from GLIMPSE v2^56^). The evaluation process was restricted to sites that had a minimum of eight reads and a genotype posterior probability of at least 0.9999. Using *GLIMPSE_concordance*, we calculated imputation errors and non-reference-discordance (NRD). NRD is defined as NRD = *(eRR + eRA + eAA) / (eRR + eRA + eAA + mRA + mAA)* where *e* and *m* correspond to the number of errors and matches, respectively, while *R* and *A* stand for reference and alternative alleles, respectively. This approach is designed to exclude the number of correctly imputed homozygous reference allele sites (*mRR*), which are the majority, therefore emphasizing the significance and giving more weight to imputation errors at alternative allele sites. To validate our results, we used the concordant dataset as validation in the first experiment, and in the second experiment, we used as validation the genotypes called with bcftools from the high-coverage non-contaminated Loschbour genome.

### g) Principal Component Analysis (PCA)

#### i) Data preparation

We generated a set of 2.8M SNPs that we used for PCA. These SNPs are a subset of the bi-allelic SNPs from the 1000 Genomes panel. We obtained this set of sites by first removing rare variants (MAF<5%). Then, we retained only sites that are included in the 1000 Genomes accessible genome strict mask, and lastly, we filtered out SNPs known to be in repeat regions. This filtering resulted in a set of 2,814,418 SNPs, which we refer to as the 2.8M SNP set.

Afterwards, we identified which positions differed based on whether genotypes were computed with bcftools or ATLAS. To do that, we intersected the imputed SNP sets from both tools so we could compare genotypes at each position, and check for agreement between the tools. Specifically, from both vcf files, we extracted the GT (genotype) columns using *bcftools query -f ‘[%GT]\n’*, we compared each position pairwise, and we determined if the imputed genotypes agreed or differed between the two sets with a python script. As for the validation samples, we similarly built validation datasets for each individual sample based on the positions concordantly identified between the two sets of genotypes called with bcftools and ATLAS on the high-coverage genomes. All SNP counts of concordant and discordant positions for both validation and imputed datasets are reported in **Table S6**.

#### ii) Performing PCA

For both PCA analyses, sample files in vcf format were first converted into the PLINK^57^ format in order to subsequently convert into the required EIGENSOFT^38,39^ format for smartPCA^38,39^. Principal components were calculated using the SGDP reference panel and all samples were projected onto the resulting components with the *lsqproject* parameter enabled. In the first experiment, we restricted the PCA to the 2.8M SNP set. The projected samples included the validation concordant samples, which served as the ground truth, as well as the sets of positions differently and concordantly imputed when genotype likelihoods were computed with either bcftools and ATLAS. In the second experiment, we restricted the PCA to the 1240K SNP set^35–37^ and projected the Loschbour ancient sample, the two present-day Dinka and Greek samples, as well as the contaminated and then imputed genomes. We projected the samples onto the SGDP reference panel, but also on a subset restricted to only the Western Eurasian populations of SGDP.

### h) Euclidean Distance

To quantify the differences between the SNP sets, we calculated an Euclidean distance between them and the validation concordant of the respective sample. This distance was calculated across the first 10 PCs, and subsequently weighted by factoring the eigenvalue of each PC.

#### i) Contamination

In order to assess the impact of contamination on imputation, we virtually contaminated an ancient European genome known as the Loschbour man^47^. Modern human DNA samples (**Table S1**), one from an African Dinka individual and another from a Greek European individual, served as the sources of contaminant DNA. We first started by downsampling the Loschbour genome to 1x coverage with samtools. Then, we computed and removed the corresponding proportion of reads from the ancient genome that would equate to the desired contamination level. As an illustration, for a contamination level of 10% with modern DNA, we initially removed 10% of the reads present in the ancient genome, then, we added an equivalent quantity of reads sourced from either the Dinka or Greek contaminant DNA.

#### j) Contamination assessment

We assessed the level of contamination using contaminationX^48^. This tool, effective for low-coverage male samples, is based on the unique property of the X-chromosome in males (hemizygous), where typically only one DNA sequence type is seen per cell. Any deviation from this, such as observing multiple alleles at a given site, indicates possible contamination, post-mortem damage, sequencing or mapping errors. We first estimated the allele counts using ANGSD^58^ with the following parameters: *-b 5000000 -c 154900000 -k 1 -m 0*.*05 -d 3 -e 20*, and specified the correct reference allele frequency panel for the sample population with the parameter *-h HapMap_pop*. Then, we estimated the level of contamination with contaminationX, using the following parameters: *maxsites=1000 nthr=4* and *oneCns=1*. We only estimated contamination up to 40% contamination rate, as beyond this, contaminationX tends to underestimate the actual contamination level as previously described^48^.

### k) Effect of contamination at the haplotype level

Before inferring local ancestry, we phased the genomes with SHAPEIT5^59^. We used the command *phase_common* while keeping rare variants (*--filter-maf 0*) and made use of the 1000 Genomes UMICH reference panel and the HapMap recombination genetic maps.

We inferred local ancestry with RFMix^60^ v2.03-r0 and used subsets of the 1000 Genomes panel as a reference. For the imputed Dinka-contaminated 1x Loschbour genomes, we had three reference populations (YRI, LWK and CEU), whereas we restricted to two reference populations (YRI and CEU) when inferring local ancestry on two African-American (labeled as ASW) genomes, NA19703 and NA20278, from the 1000 Genomes panel.

#### l) Text quality enhancing through the use of specialized AI tools

We used AI tools, specifically ChatGPT, to improve the quality of the text in the introduction section of our article.

## Data availability

The eight ancient individual samples analyzed in this study are publicly available and were first described in the following studies: ela01^45^ (https://doi.org/10.1126/science.aao6266); BOT2016 & Yamnaya^42^ (https://doi.org/10.1126/science.aar7711); NE1^41^ (https://doi.org/10.1038/ncomms6257); SZ1^46^ (https://doi.org/10.1038/s41467-018-06024-4); Sumidouro5^43^ (https://doi.org/10.1126/science.aav2621); Yana^44^ (https://doi.org/10.1038/s41586-019-1279-z); and Loschbour^47^ (https://doi.org/10.1038/nature13673). The SGDP reference panel^40^ dataset in plink format can be found in https://sharehost.hms.harvard.edu/genetics/reich_lab/sgdp/variant_set/. We downloaded from Seven Bridges Cancer Genomics Cloud the bam files aligned to hg19 reference genome for two SGDP genomes, Dinka-3 and Greek-2. The 1000 Genomes v5 phase 3 panel resequenced to 30X coverage is available at the European Nucleotide Archive under accession code ERP114329 (https://www.ebi.ac.uk/ena/browser/view/PRJEB31736).

## Acknowledgements

We want to thank Dilek Koptekin, Théo Cavinato and Samuel Neuenschwander for fruitful discussions. A.G.M. and B.S.d.M. were supported by a Swiss National Science Foundation (SNSF) project grant (PP00P3_176977) to O.D., and by an SFNS grant (PCEGP3_181251) and a European Research Council grant (CAMERA 679330) to A.-S.M. S.R. was funded by an SNSF project grant (PP00P3_176977) to O.D. and by an SNSF Postdoc.Mobility project grant (P500PB_211106).

## Author’s contributions

B.S.d.M., S.R. and O.D. conceptualized and designed the study. A.G.M. and B.S.d.M. conducted the experiments. A.G.M. and B.S.d.M. wrote the manuscript with input from O.D, A.-S.M and S.R. This work has been supervised by B.S.d.M. All authors interpreted the results and reviewed the final manuscript.

## Code availability

The scripts used to generate the data and results presented in our study are provided in this github repository. https://github.com/TozeMarques/PMD_Contamination_Impact_on_aDNA_Imputation

