## Supplementary Information file for "Assessing the impact of post-mortem damage and contamination on imputation performance in ancient DNA"

### Supplementary Note 1: Post-mortem damage patterns

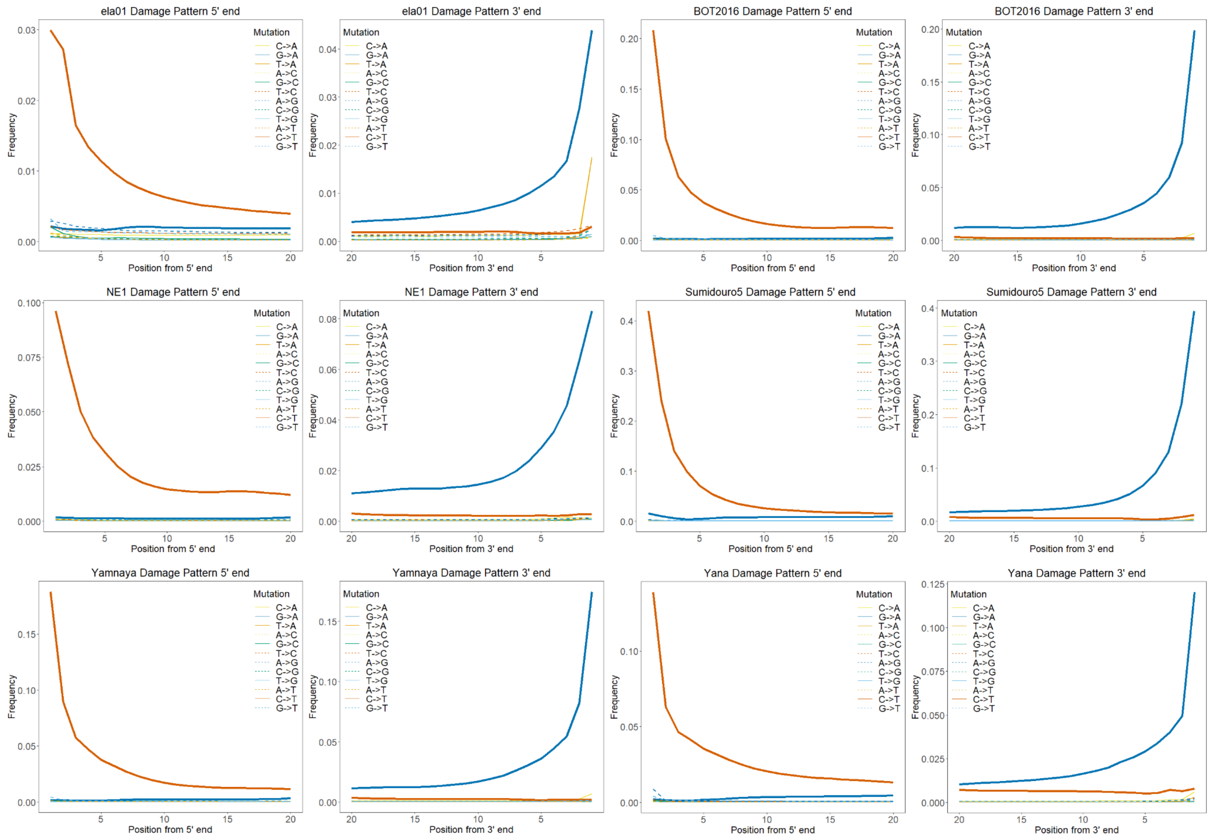


**Supplementary Figure 1: Substitution rates at the ends of the reads (5' and 3' ends).** Frequency of nucleotide substitutions at the 5' and 3' ends of sequencing reads measured with the bamdamage tool from bammds. Nucleotide substitution rates are presented for six of the studied ancient individual samples (ela01, BOT2016, NE1, Sumidouro5, Yamnaya, Yana). Each panel shows the substitution frequency on the *y*-axis in function the position on the first 20 nucleotides on the *x*-axis.

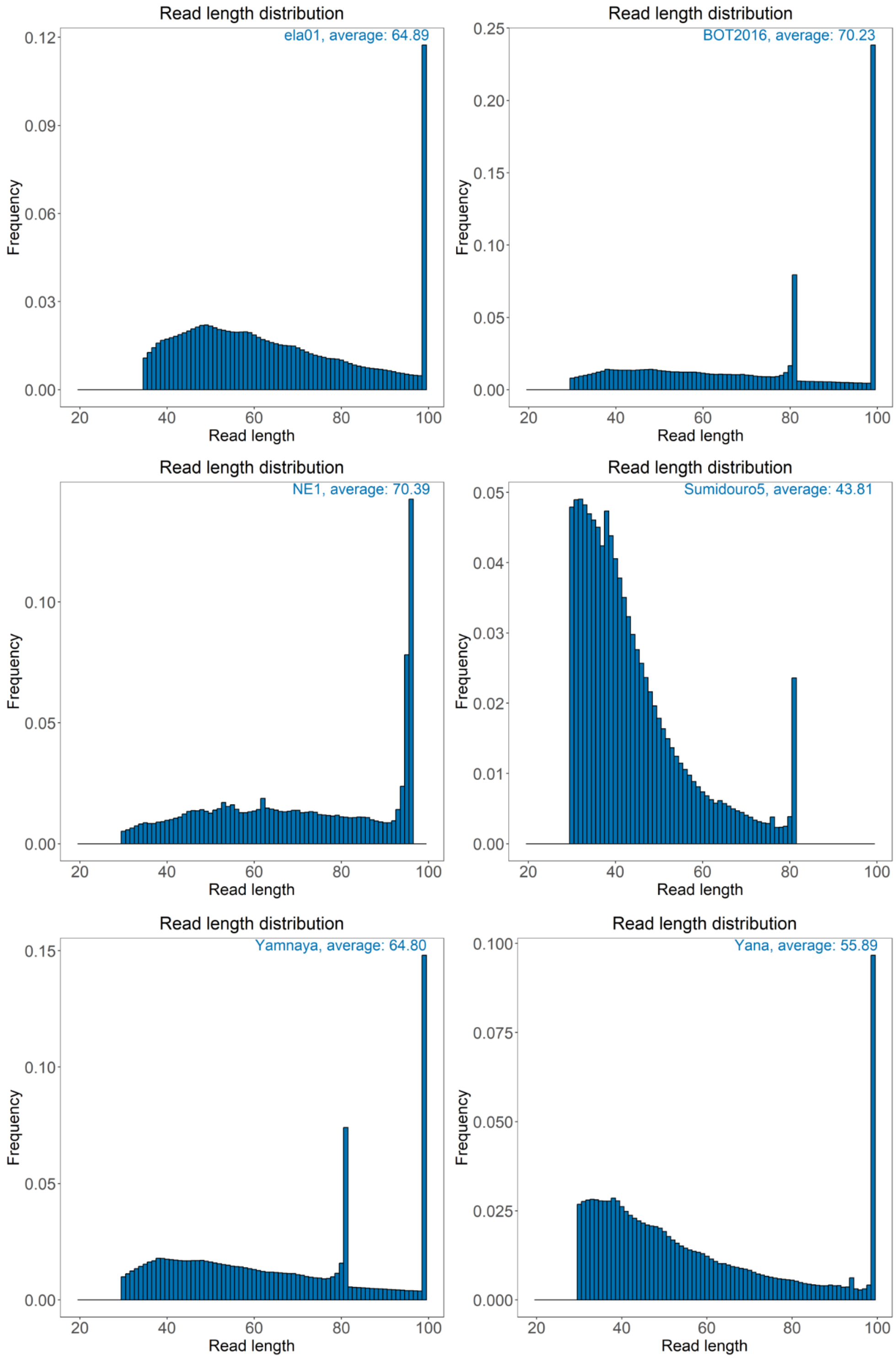


**Supplementary Figure 2: Read length distribution.** Distribution of the read length for six of the studied ancient individual samples (ela01, BOT2016, NE1, Sumidouro5, Yamnaya, Yana). The histograms represent the frequency of reads on the *y*-axis of a certain read length on the *x*-axis, for each sample. The average read lengths are denoted on the top right of each panel.

### Supplementary Note 2: Enhanced performance of ATLAS in high-coverage genotype calling

Establishing an accurate ground truth for ancient genomes, even when high-coverage genotypes are available, is a challenge due to PMD prevalence. In this study, we found PCA to be a useful evaluation framework to compare differently generated datasets. For the high-coverage genomes, we observed that positions differently called by ATLAS and bcftools (see methods for more details on quality control procedures) display distinct spatial patterns in the PCA space (**Supplementary Figure 3**). Specifically, genotypes differently identified with ATLAS consistently positioned closer to the validation set. In contrast, those identified by bcftools were more distant, with the exception of the individual sample ela01. However, these discrepancies can be partly explained by the difference in the number of SNPs between the two as the bcftools discordant SNP set contained fewer SNPs, averaging around 10,500, resulting in an important variation in the number of SNPs across the samples when compared with the ATLAS discordant SNP set that averages approximately 472,000 SNPs (**Table S6**).

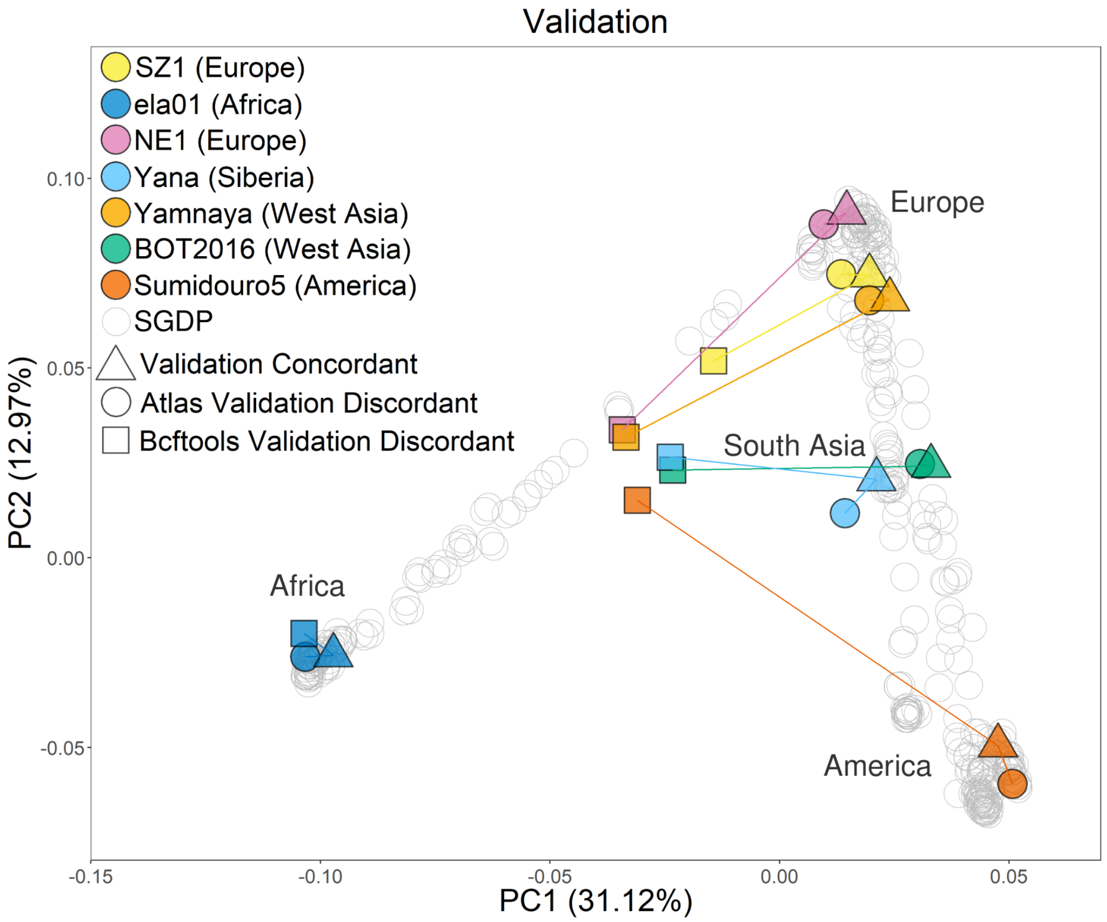


**Supplementary Figure 3: PCA Validation.** Two first principal components of principal component analysis (PCA) of present-day genomes (SGDP). The genetic data of seven ancient individuals using the 2.8M SNP set were projected onto it. For each individual, the triangle shape indicates the validation concordant dataset, and the circles and square shapes the positions differently identified by ATLAS ("ATLAS Imputed Discordant") and bcftools ("Bcftools Imputed Discordant"), respectively.

### Supplementary Note 3: Computational resources usage by ATLAS and bcftools to generate genotype likelihoods

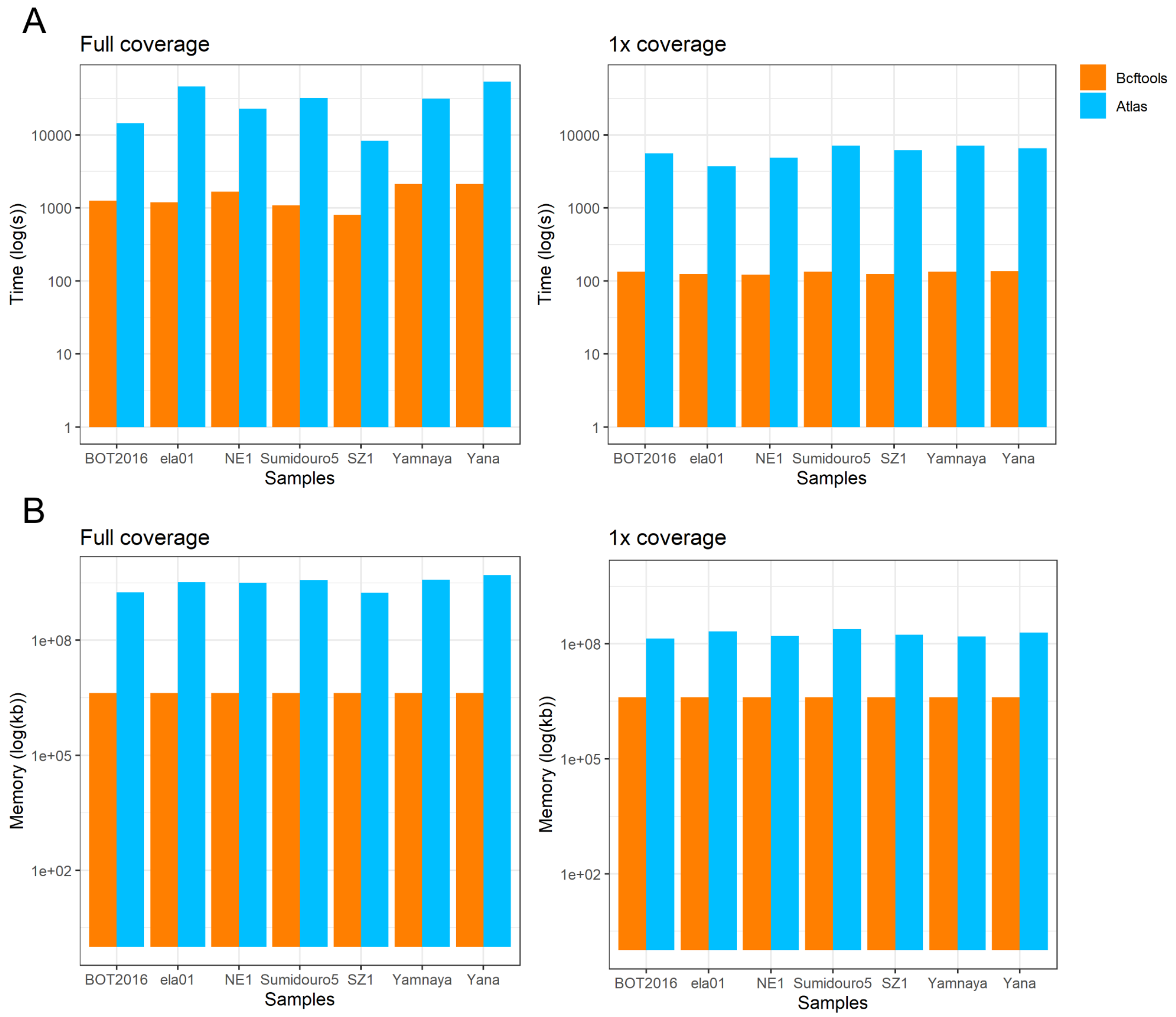


**Supplementary Figure 4: Run time and computational resources when using ATLAS and bcftools**. Disparity in total run time (**A**) and memory usage (**B**) between ATLAS (blue) and bcftools (orange) during the genotype calling process of seven ancient individuals WGS. This comparison was performed on both high-coverage genomes (left plots) and downsampled 1x coverage genomes (right plots). Run time was measured on chromosome 1, while memory usage was assessed on the 22 autosomes. Results shown (*y*-axis) are represented in a log10 scale.

### Supplementary Note 4: Post-mortem damage impact on imputation

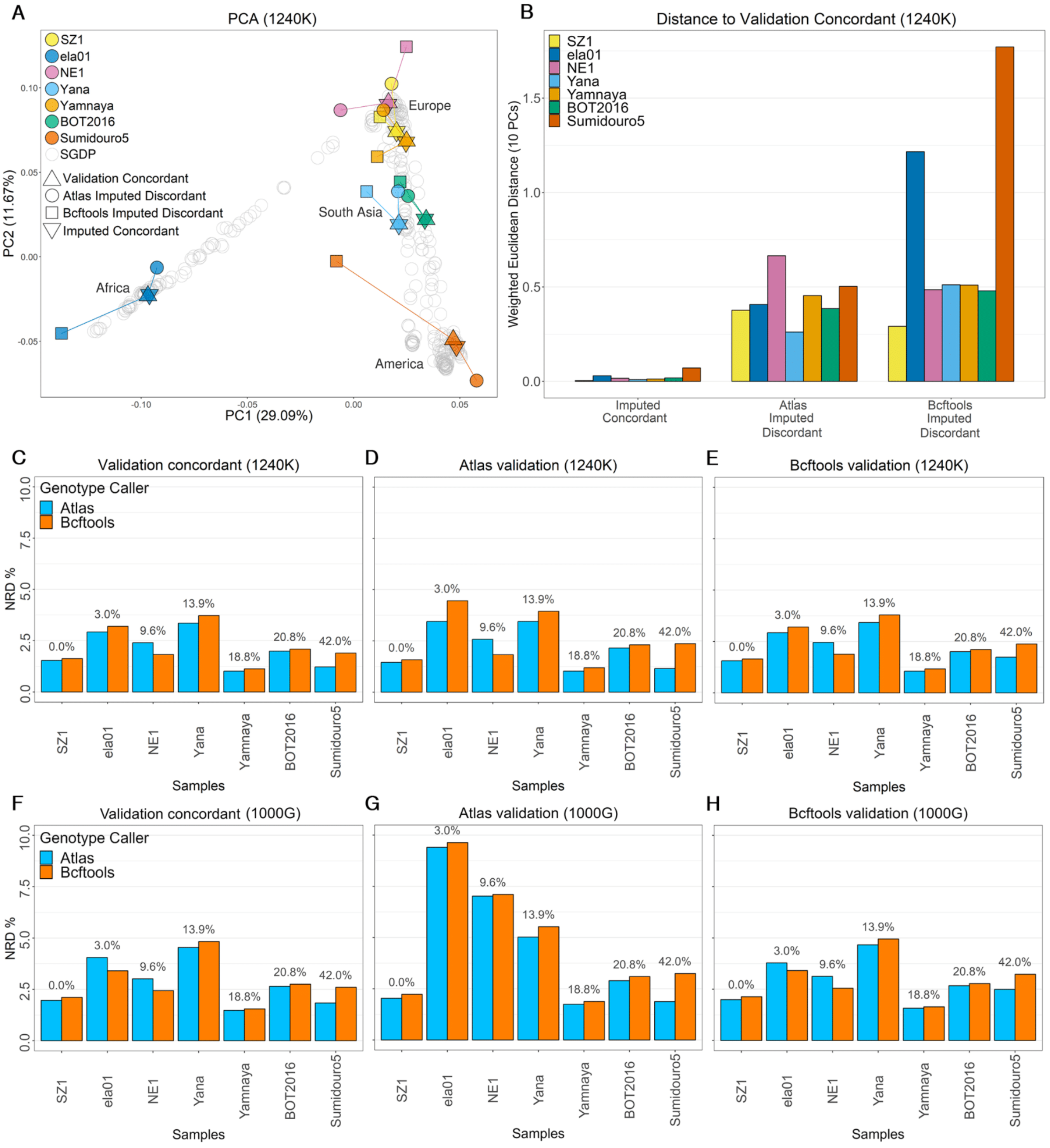


**Supplementary Figure 5: Effect of two different genotype callers and SNP sets on imputation accuracy. A**) Two first principal components of principal component analysis (PCA) of present-day genomes (SGDP) with the 1240k SNP set. The genetic data of seven ancient individuals were projected onto it. The triangle shape indicates the validation concordant dataset, the inverse triangle the set of imputed positions in agreement with both tools, and the circles and square shapes the positions differently identified by ATLAS ("ATLAS Imputed Discordant") and bcftools ("Bcftools Imputed Discordant"), respectively. **B**) Weighted Euclidean distances for the seven ancient individuals with the 1240k SNP set. Euclidean distances between the validation concordant set of each sample, and their concordant and discordant imputed genotypes were calculated across the first 10 PCs, with distances weighted by the eigenvalue of each PC. For additional information on the concordant and discordant samples SNP counts, refer to **Table S7. C**) Non-reference discordance (NRD) for the seven ancient individual samples with the 1240k SNP set when called with either ATLAS (blue) or bcftools (orange) prior to imputation. The values on top of each sample indicate the PMD rate. NRD was assessed using three validation datasets: (**C**) validation concordant, (**D**) ATLAS validation, and (**E**) bcftools validation. **F**) Assessment of NRD for the seven ancient individual samples with the 1000 Genomes (1000G) SNP set when called with either ATLAS or bcftools. The PMD rate is shown on top of each sample. The NRD was calculated using the: (**F**) validation concordant, (**G**) ATLAS validation, and (**H**) bcftools validation datasets.

### Supplementary Note 5: Contamination and imputation

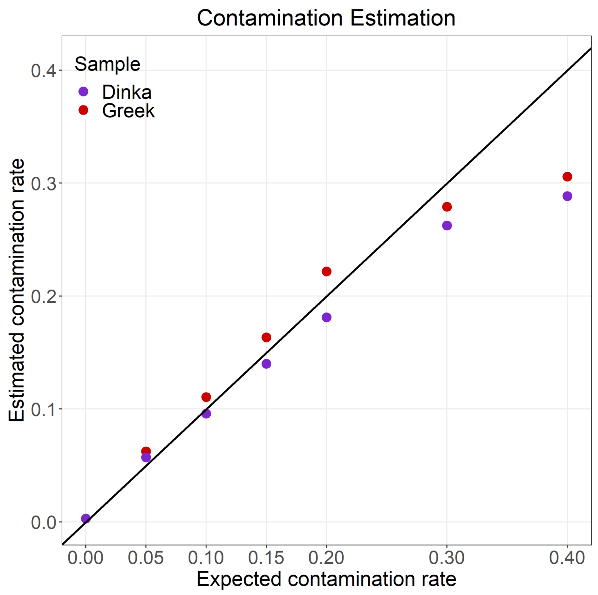


**Supplementary Figure 6: Contamination estimation on chromosome X**. Comparison of estimated (*y*-axis) with expected (*x*-axis) contamination rates when introducing different amounts of reads from modern human genomes in the Loschbour downsampled (1x) genome. Two contaminants were used: a present-day Greek genome in red, and a present-day Dinka genome in purple. The black line indicates equality between expected and estimated contamination (*y*=*x*).

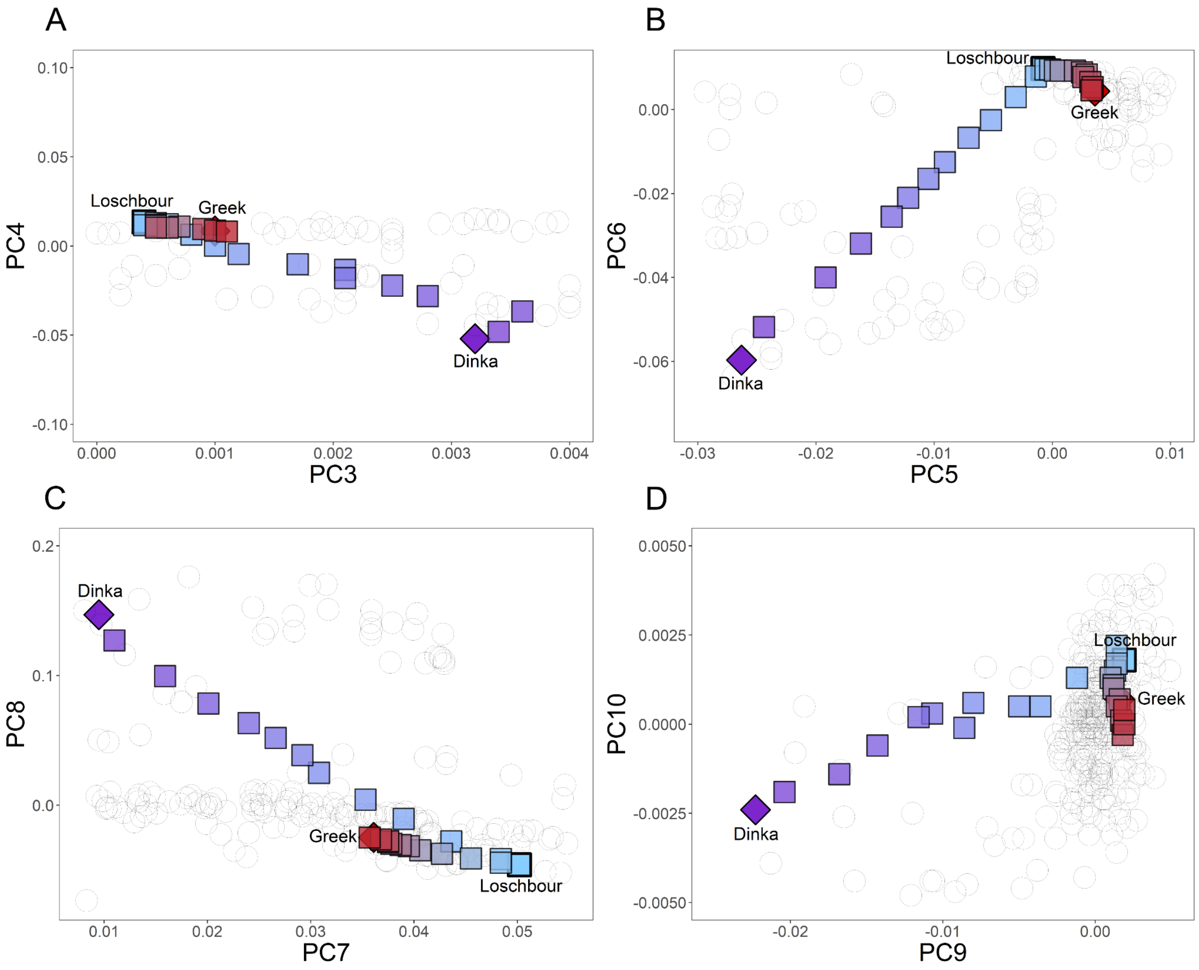


**Supplementary Figure 7: Effect of contamination on genotype imputation on higher PCs.** Principal component analysis (PCA) of present-day genomes, restricted to 1240k SNPs, and projection of uncontaminated and imputed contaminated ancient genomes. The ancient downsampled (1x) Loschbour genome (light blue, in bold) was subjected to varying degrees of contamination with DNA from a present-day Greek individual (red, in bold) and a present-day Dinka individual (purple, in bold). All imputed contaminated downsampled Loschbour genomes are projected on top of SGDP populations (gray circles). The resultant projections are displayed across **A)** PC3 vs PC4, **B)** PC5 vs PC6, **C)** PC7 vs PC8 and **D)** PC9 vs PC10.

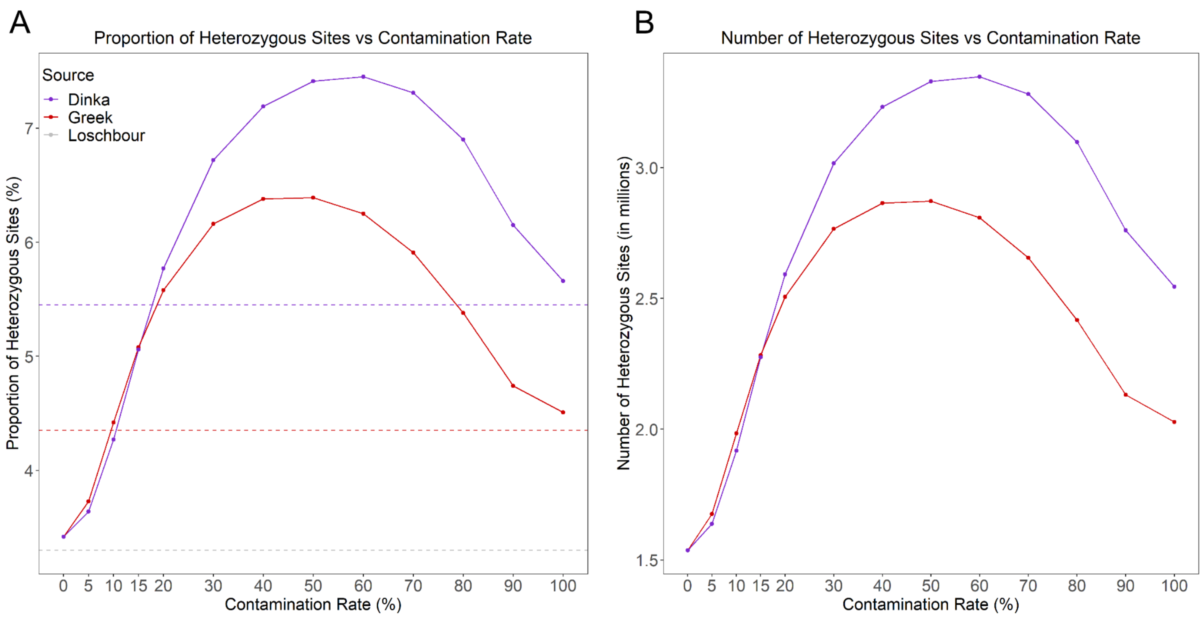


**Supplementary Figure 8: Heterozygous positions across different contamination rates.** Quantification of heterozygous positions across varying contamination rates ranging from 0% (imputed 1x Loschbour genome) to 100% (imputed 1x Greek or Dinka genomes) contamination. **A)** Proportion of heterozygous sites for Dinka-contaminated Loschbour genomes in purple, and Greek-contaminated genomes in red. Dashed lines represent the proportion of heterozygous sites for the high-coverage genomes. **B)** Absolute number of heterozygous sites (in millions) for Dinka- and Greek-contaminated genomes.

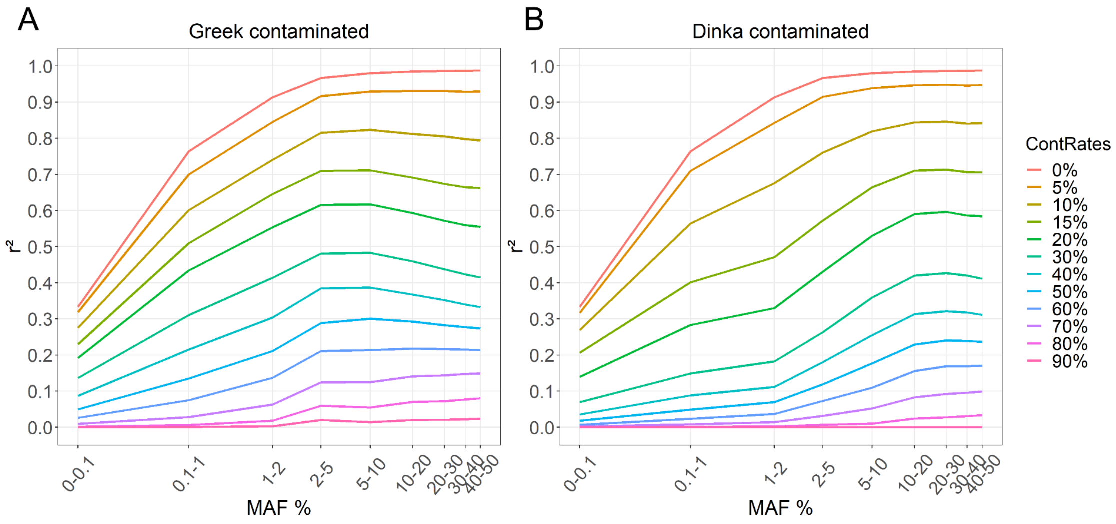


**Supplementary Figure 9: Impact of contamination on genotype imputation**. Mean imputation accuracy ($r^2$) as a function of minor allele frequency (MAF) of the downsampled contaminated Loschbour genome ranging from 0% to 90% contamination. Loschbour genome was downsampled to 1x and contaminated with **A)** present-day Greek DNA, and **B)** present-day Dinka DNA.

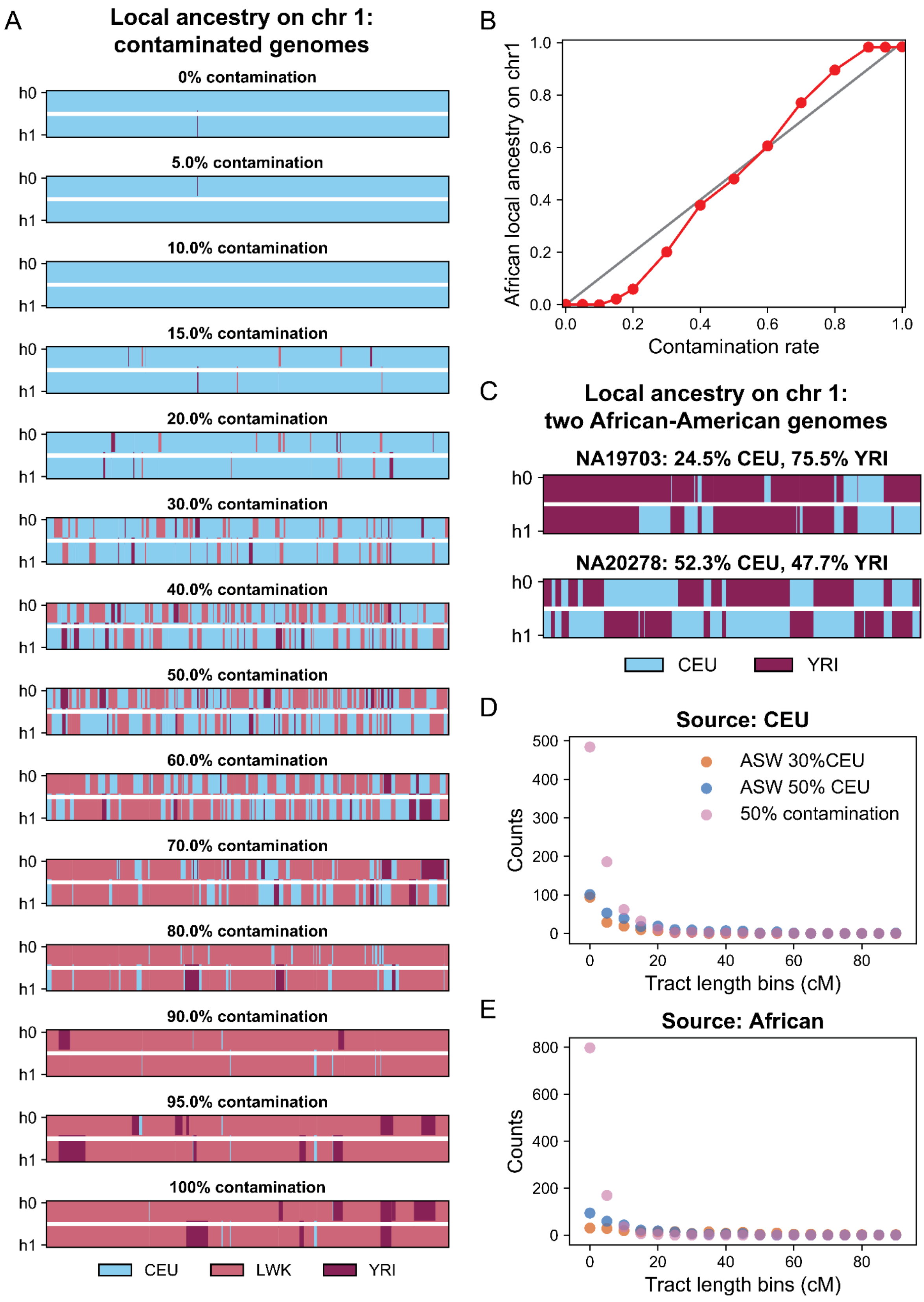


**Supplementary Figure 10: Local ancestry inference for contaminated imputed chromosomes. A)** Local ancestry inferred on chromosome 1 for contamination rates varying from 0% (imputed 1x Loschbour genomes) to 100% (imputed 1x Dinka genome) using three reference ancestries: CEU (Utah residents with Northern and Western European ancestry), LWK (Luhya in Webuye, Kenya) and YRI (Yoruba in Ibadan, Nigeria). **B)** Ratio of African (LWK+YRI) local ancestry on chromosome 1 as a function of contamination rate. **C)** Inferred local ancestry on chromosome 1 for two African American individuals, NA19703 and NA20278, in the 1000 Genomes panel (ASW population label) using two reference ancestries: CEU and YRI. **D)** Distribution of CEU-inferred tracts on the African American genomes and on the contaminated Loschbour genome (50% contamination with reads from a present-day Dinka). **E)** Distribution of African-inferred tracts on the African American genomes and on the contaminated Loschbour genome (50% contamination with reads from a present-day Dinka). For the contaminated genome, African-inferred tracts include LWK and YRI tracts.
